# Is critical brain dynamics more prevalent than previously thought?

**DOI:** 10.1101/2025.09.02.673722

**Authors:** Antonio J. Fontenele, J. Samuel Sooter, Ehsan Ziarati, Andrea K. Barreiro, Cheng Ly, Woodrow L. Shew

## Abstract

The hypothesis that the brain operates near criticality has far-reaching implications for brain function and is supported by growing experimental evidence. Observations of scale-invariant brain activity agree with this hypothesis, but what about when brain activity is not scale-invariant? Should we reject the criticality hypothesis When power-laws poorly fit the data or when strong oscillations occur (dominated by a specific time scale)? Here we show several ways that criticality can be hidden from traditional data analytic approaches, leading to false negative conclusions. We use a parsimonious high-dimensional model to demonstrate how neural systems may separate different dynamical modes into different subspaces, simultaneously generating non-critical dynamics, critical oscillations, and scale-invariant avalanches. Our results point to a need for new methods capable of revealing hidden criticality and suggest that criticality could be more prevalent than previously thought, hidden in subspaces not readily revealed by standard data analyses.

Does the brain operate in a dynamical regime close to a critical phase transition? This question is fundamental; the nature of neural computation is strongly impacted by whether a system is close to or far from criticality [1–5]. Evidence suggesting that the brain operates close to criticality has accumulated at an accelerating pace over the past two decades (a recent meta analysis found 31 reports in 2024 alone [2]). The two most common types of experimental evidence have been based on neuronal avalanche analysis [1, 6, 7] and long-range temporal correlation analysis [8–12]. Both these approaches examine a one-dimensional time series of collective neural activity, averaged over large populations of neurons, seeking temporally-correlated, scale-invariant fluctuations that obey specific scaling laws predicted by theory. In many cases, measurements agree with these predictions; the success and persistence of the criticality hypothesis rests on these cases. However, it is not difficult to find measurements that do not agree, with fluctuations poorly described by power-laws and scaling laws. Another common observation that, naively, seems to contradict criticality is oscillatory brain activity with a particular dominant frequency. At first glance, either the lack of power-laws or the prominence of a dominant oscillatory time scale appear to contradict the criticality hypothesis.

What should we conclude from these apparently contradictory observations? Assuming that all the experimental observations are sound and valid, there are two possible explanations, not necessarily mutually exclusive. First, it could be that there are biophysical control parameters constantly in flux in the brain, causing shifts in proximity to criticality. In this scenario, non-scale-invariant data reflect true deviations from criticality and we should conclude that the brain is, at times, closer to and, at other times, further from criticality. This view has been proposed in multiple previous studies and is consistent with substantial experimental evidence [6, 13–19]. However, here we argue that there is an important, but rarely considered, second plausible explanation. It could be that traditional methods for assessing criticality can be fooled. In this case, the system dynamics could be truly critical and scale-invariant, but not visible to traditional methods. We first show why such hidden criticality is plausible considering general mathematical arguments and relevant experimental evidence. Then, we use a parsimonious computational model to demonstrate concrete examples of ground truth scale-invariant critical dynamics that are missed by traditional analyses of population average activity. Moreover, our model shows how critical oscillations, avalanches, and other non-critical modes of dynamics can coexist, by separating each different mode into a distinct low-dimensional subspace. All together, our results suggest that the combination of these various subspaces make up the high-dimensional dynamical system commonly observed in the brain. Our results reconcile multiple, seemingly discrepant, experimental observations and describe prospects for new methods that can reveal hidden criticality.

## I. HIDDEN CRITICALITY

Criticality is defined by two necessary and sufficient conditions; a system is at criticality if and only if it has 1) scale-invariant behavior and 2) is at a boundary in parameter space. Given some measured data, for instance, the spike times of a large population of neurons, how have previous studies sought evidence for criticality? The vast majority of cases (more than 80% according to a recent meta-analysis [2]) begin by reducing the population activity to a one-dimensional population average (or sum) time series. Moreover, many other collective signals like EEG, MEG, and fMRI, might be reasonably approximated as a population average of underlying neural activity. With the population average time series in hand, the next step is to seek evidence for scale-invariant fluctuations in the population average time series. Most previous studies assume that if the brain is near criticality, then we will see scale-invariant fluctuations in the population average time series. More specifically, scale-invariance is sought with avalanche analysis, long-range temporal correlation (LRTC) analysis, estimates of branching ratios, and temporal renormalization group methods. In this section, we show that this assumption is sometimes correct (Case 1 below), but can be wrong in many realistic scenarios (Cases 2 and 3 in Fig. 1 and below); criticality can be hidden from view when analyzing the population average. The most important conclusion of our work here is that criticality may be far more prevalent that previously thought because criticality is easy to hide from the most common analyses.

**FIG. 1.**
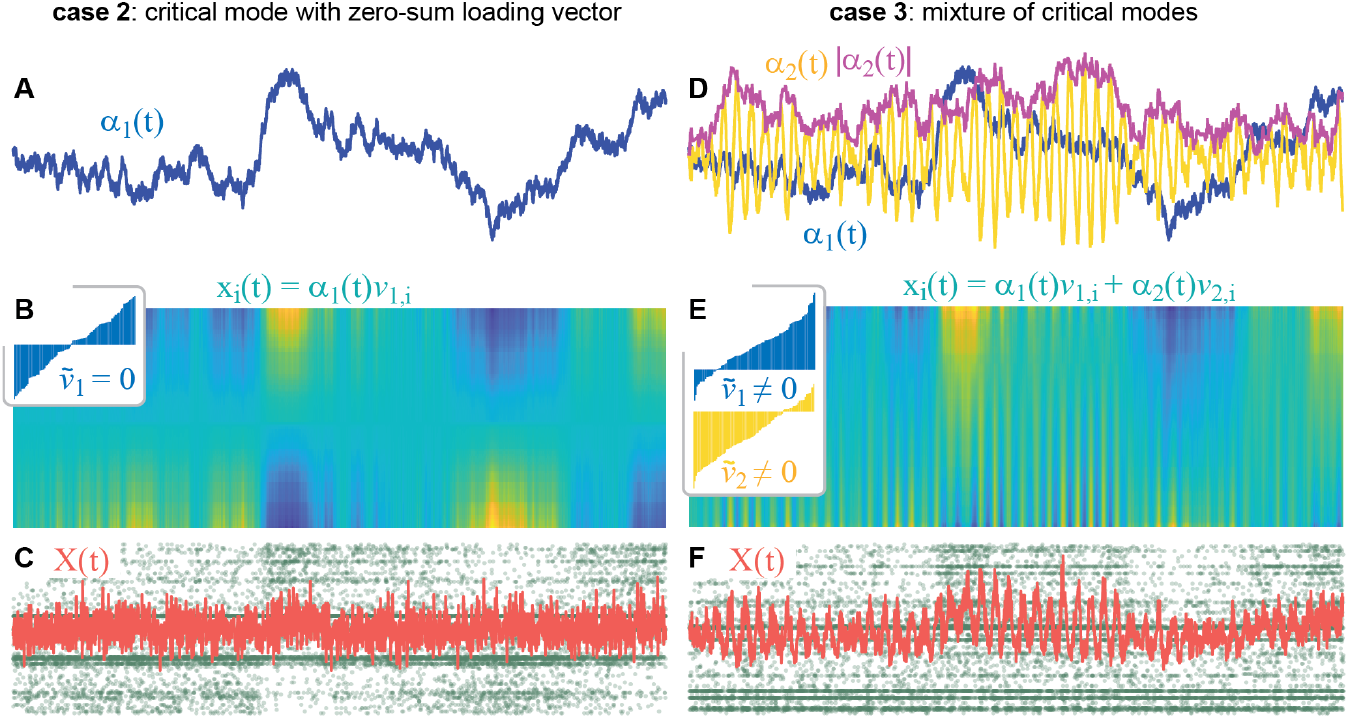
Two cases of hidden criticality. **A** Coefficient time series *α*(*t*) for a mode very near criticality (*λ* = 0.99). **B** For a basis vector with near zero component sum (inset shows vector components for 100 neurons), the population activity (color) will have strongly anti-correlated groups of neurons. Each row of the colored activity raster represents one neuron’s activity time series. **C** The population average activity (red) does not reveal the Critical dynamics *α*_1_(*t*) that are clear in panel A and in the spike raster (green), because anticorrelated activity cancels out in the population average. **D** Coefficient time series for two critical modes, one oscillatory with *λ* = 0.99 + 0.3*i* (yellow), the other identical to that in panel A (blue). Note that the amplitude (magenta) of the critical oscillatory mode is scale invariant as in eq. 13. **E** We consider basis vectors for each mode with non-zero component sum. **F** Mixing the two modes in the population average (red) hides the underlying critical dynamics of *α*_1_(*t*) and *α*_2_(*t*). For cases 2 and 3, traditional criticality analyses would arrive at false-negative conclusions about criticality. The single mode dynamics shown in panels A and D were simulated according to eq. 7 and then used to construct the population activity in panels B, C, E, and F (see Methods).

To demonstrate this concretely, in this section, we examine some basic mathematical properties of population average time series. In the next section, we demonstrate several specific cases in more detail using a commonly-studied computational model. We begin by noting that the activity 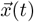 of *N* neurons can always be decomposed as a linear combination of *N* modes using basis vectors 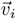,

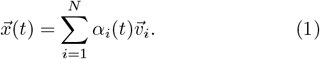

Then the population average time series *X*(*t*) can be written as

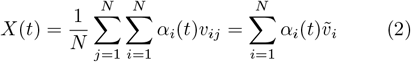

where 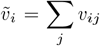 is component sum of 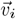. In the next section, we will use the eigenvectors of an interaction matrix as the basis, but more generally, this can be done with any set of vectors that span *R*^*N*^ . Common ways to find such basis vectors include principal component analysis (PCA), canonical correlation analysis (CCA), shared-variance component analysis (SVCA), and many other dimensionality reduction methods. Not all modes contribute equally to the population average; one mode could dominate or a mixture of modes could contribute. Eq. 2 makes clear that the contribution of mode *i* is determined by two key factors: 1) the variance of the co-efficients *α*_*i*_(*t*) and 2) the sum of the corresponding basis vector. If 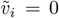, the mode will be invisible in the population sum (zero contribution to the sum in eq. 2), regardless of the variance of *α*_*i*_(*t*). More generally, if 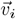 has positive and negative entries that cancel, then 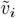 will be small. In real data, this scenario occurs when the activity of some neurons is anticorrelated with that of other neurons. Such negative covariances among neurons is often observed in experiments [20–26]. Thus, *α*_*i*_(*t*) could be temporally scale invariant (i.e. critical), but could be missed by any analysis of the population average if 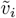 is sufficiently small. This scenario is illustrated in Fig. 1A-C. Below we will show that scenarios like this arise easily in our model and many previous models (especially in the context of working memory) and are consistent with many experiments. Most notably, Dahmen and colleagues studied this phenomenon and referred to it as a “second type of criticality” [21]. Below, we will discuss further whether it is really another type of criticality or just the usual type, but hidden. Nonetheless, it is clear that critical dynamics can be missed when analyzing the population average. A more general description of this scenario is that any dynamics that live within the *N* − 1 dimensional subspace that is orthogonal to the all-ones vector 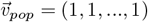 will be invisible to analyses based on the population sum. Indeed, all vectors that are orthogonal to 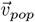 have zero component sum.

A second important situation in which traditional analyses of the population average may arrive at false negative conclusions about criticality is when many independent modes are mixed together, especially when one or more dominant modes are oscillatory. This can occur when many terms contribute significantly to the sum in eq. 2. A simple example with two near-critical mixed modes is illustrated in Fig. 1D-F. With the mixture of modes, it can be difficult to extract the underlying critical dynamics. For instance, critical oscillatory modes can confound traditional neuronal avalanche analysis and non-oscillatory critical modes can confound traditional LRTC analysis.

## II. LINEAR RATE MODEL

Next we demonstrate how criticality can be hidden in a commonly studied model of neuronal population dynamics [21, 27–30]. We consider a simple model that describes the firing rates of *N* interacting neurons. We will first review what criticality means in this model. Then we will show in detail several example cases: Case 1 when traditional assumptions hold and population average signals reveal criticality without issues, Case 2 - when criticality is hidden due to anticorrelated neurons, Case 3 - when criticality is hidden due to mixed modes. As we proceed, we will discuss how these examples offer plausible explanations of some experimental findings that were thought to be discrepant.

The model is

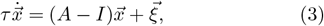

where the activity of the *i*^*th*^ neuron at time *t* is the *i*^*th*^ entry of 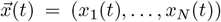. The noise 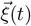 is uncorrelated Gaussian white noise, i.e., 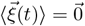 and ⟨*ξ*_*i*_(*t*)*ξ*_*j*_(*t*^*′*^)⟩ = *σ*^2^*δ*_*ij*_*δ*(*t* − *t*^*′*^). The entry *A*_*ij*_ of the interaction matrix represents the interaction from neuron *j* to neuron *i*; *A* may be asymmetric and may have some negative columns to represent inhibitory neurons. The matrix *A* has right eigenvectors 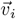 with eigenvalues *λ*_*i*_ and left eigenvectors 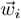. In general, the right eigenvectors are not mutually orthogonal, but the left and right eigenvectors are biorthogonal. If the eigenvectors span *R*^*N*^, as in all the cases we examine below, the population activity vector can be decomposed into a linear combination of *N* modes as in eq. 1, as can the noise,

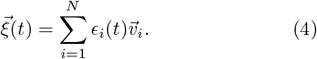

Substituting eqs. 1 and 4 into eq. 3, we get

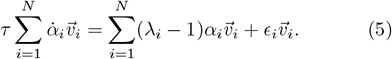

Then, multiplying by a particular left eigenvector, we have

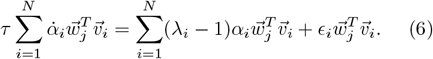

Since 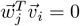 for all *i* ≠ *j*, all terms in the sums are zero except the *j*^*th*^. Thus, each mode of the system evolves independently, according to

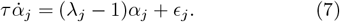

Stochastic calculus affords the following solution to this equation (dropping the *j* subscript),

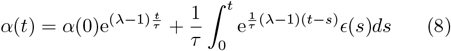

For a complex eigenvalue, *λ* = *k* + *ib*, the solution is

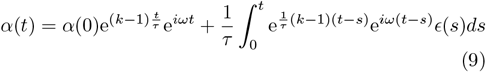

where *ω* = *b/τ*. Whether a mode *α*(*t*) exhibits critical dynamics or not is determined by *k*, i.e. the real part of that mode’s eigenvalue. As *k* decreases towards extreme negative values, exp((*k* − 1)*t/τ*) approaches zero. And, the factor in the integrand exp((*k* − 1)(*t* − *s*)*/τ*) approaches zero unless *s* = *t*, in which case it is 1. Thus, for extreme negative *k, α*(*t*) approaches white noise

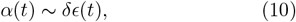

where the noise amplitude *δ* decreases as *k* decreases. If *k >* 1, the dynamics are unstable; *α*(*t*) blows up to *±* infinity. A mode with *k* = 1 meets both defining criteria for criticality: 1) it is at a boundary in parameter space and 2) it has scale-invariant fluctuations [2]. Here the control parameter is *k* and the relevant boundary in parameter space is at *k* = 1, the edge of instability. To see how the dynamics are scale-invariant at this boundary, consider first the case with a purely real eigenvalue (*b* = 0). The solution becomes

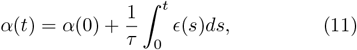

which is a one-dimensional random walk in the continuum limit. A random walk is temporally scale-invariant with known critical exponents and scaling laws [31, 32]. If a mode has a complex eigenvalue with *k* = 1, we have

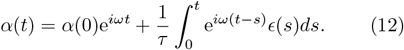

Note that, in this case, *ϵ*(*s*) is complex Gaussian noise, because the corresponding eigenvector in eq. 4 is complex. Rewriting this, we see that we have an oscillation with frequency *ω* whose amplitude is modulated according to a random walk

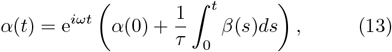

where *β*(*s*) = e^−*iωs*^*ϵ*(*s*) is also complex Gaussian noise with the same statistics as *ϵ*(*s*). Thus, for a critical complex mode, *α*(*t*) exhibits oscillations with a particular frequency, i.e. is not scale invariant, but the amplitude of the oscillation |*α*(*t*) | is scale-invariant. This observation is reminiscent of traditional studies of long-range temporal correlations (LRTC). In this tradition, a measured brain signal (often MEG or EEG) is filtered to isolate a particular frequency band and then the oscillation amplitude envelope time series is analyzed, seeking temporal scale invariance [8–11].

Put simply, criticality is fundamentally lowdimensional in this model. If only one mode has *k* near 1, then criticality is 1-dimensional. This situation is easy to instantiate in our model as we will show below (Case 1). If *m* modes have *k* near 1, then we should conclude that the system has a *m*-dimensional critical subspace (Case 3 below and similar to claims from experiments in mouse motor cortex [7]). If some of these near critical modes are complex, we can easily end up with a situation with coexistent oscillations and critical dynamics and a messy population sum from which it is difficult to discern scale-invariance. We will also demonstrate this scenario with our model below (Case 3). It is important to realize however, that while modes evolve independently, neurons do not. Many (typically all) neurons participate in each independent mode. Thus, if many modes are near criticality (Case 3), we expect broadly distributed pairwise covariances across neurons, as investigated in detail in multiple recent studies [21, 27–29, 33].

## III. DEMONSTRATING TYPES OF HIDDEN CRITICALITY

We now consider how this model behaves for three example interaction matrices *A*, which clearly demonstrate how critical dynamics can be hidden or apparent in the population average dynamics. For each case, we simulate the full *N*-dimensional dynamics and then isolate particular modes to reveal criticality and/or critical oscillations. For a given multivariate time series 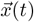 generated by our model (eq. 3, Methods), we isolate the dynamics *α*_*j*_(*t*) of mode *j* by multiplying by the left eigenvector 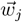 and normalizing by the inner product 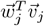,

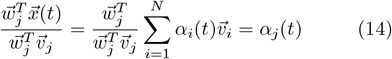

For modes with a complex eigenvalue, *α*(*t*) is complex. For these modes we take the magnitude |*α*(*t*) | to obtain the amplitude envelope of the oscillations. For each isolated mode, we perform the most common tests for assessing criticality, analyzing *α*(*t*) for real modes and |*α*(*t*) | for complex modes. This includes avalanche analysis (size and duration distributions and crackling noise scaling), DFA analysis, power spectra, autocorrelation functions, and a relatively new measure of distance from criticality *d*_2_ based on a temporal renormalization group approach [34]. For avalanche analysis, power spectra, and DFA, a larger range of power-law scaling should be interpreted as evidence for nearer proximity to criticality; as a system moves away from criticality the power-law range decreases.

### Case 1. Non-hidden, non-oscillatory criticality

First, we demonstrate an example when critical dynamics are not hidden. In this case, the population average reveals criticality perfectly well, which is consistent with many previous observations of scale invariance using traditional analyses of population average signals (some examples include refs. [1, 7, 13, 35], see ref. [2] for a systematic review). We define the interaction matrix similar to previous studies of probabilistic binary neuron models [36, 37]. The entries of *A* are first drawn from a uniform distribution [0, 1] and then 20% of columns are multiplied by -1 to model inhibitory neurons. Finally, we ensure that the dominant mode is close to criticality by multiplying the entire matrix by a constant chosen to set the real part of its largest eigenvalue to *k* = 0.999. The result is that the dominant mode has a purely real eigenvalue while all other modes have eigenvalues around zero (Fig 2B). The dominant mode has the largest fluctuations and large eigenvector component sum (Fig 2C). Thus, the near-critical dominant mode is the only mode with significant impact on the population average signal (Fig 2D,E) and all the traditional assays produce strong evidence for criticality (Fig 2F-K) - large power-law ranges for avalanches and the power spectrum, and DFA exponents well above 0.5.

**FIG. 2.**
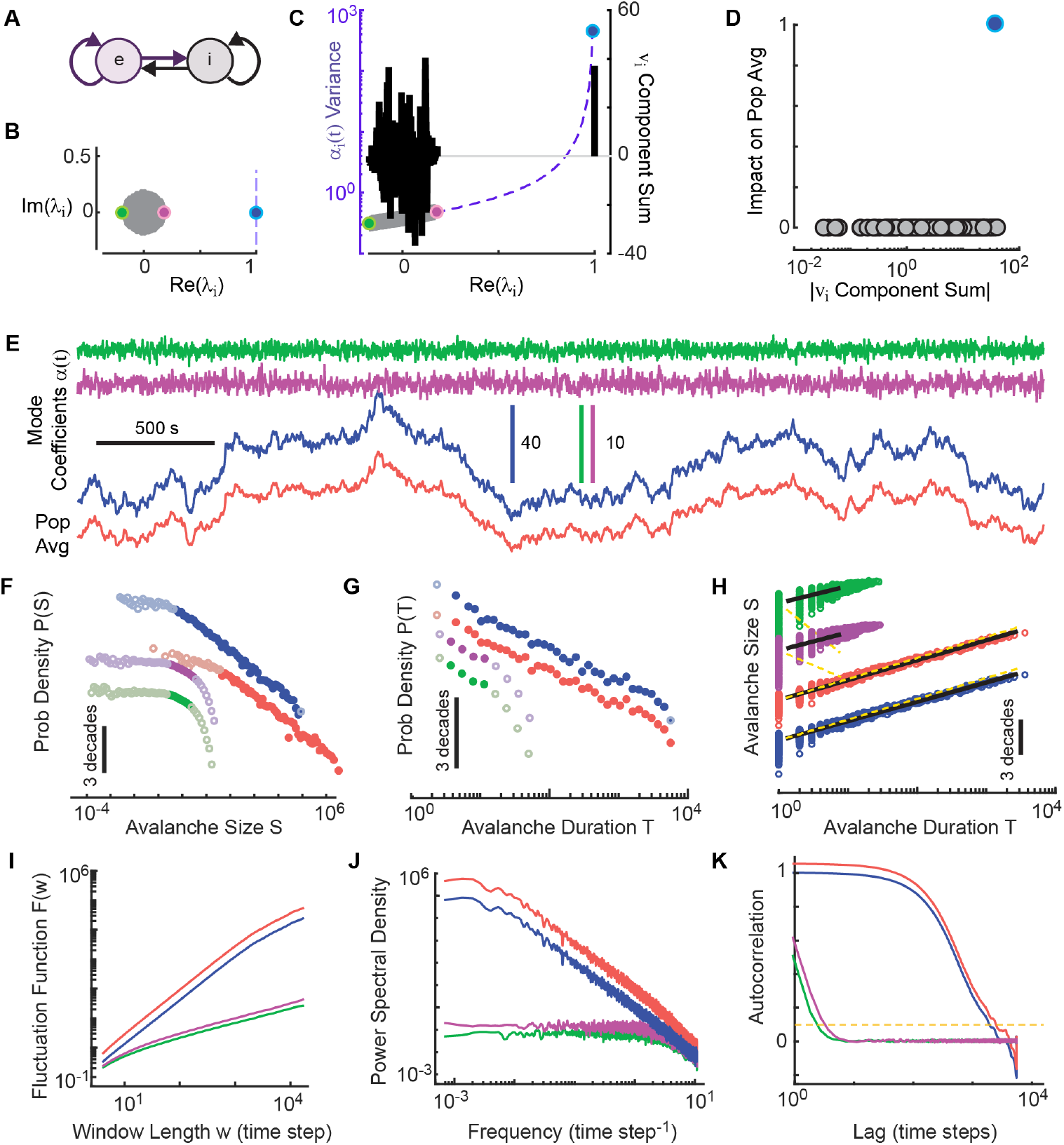
Case 1: Non-hidden, non-oscillatory critical dynamics. **A** Schematic of interaction matrix *A* with a group of excitatory neurons (e) that interacts with a group of inhibitory neurons (i). Dale’s law is enforced. **B** Bulk of eigenvalues of A are near zero (gray, magenta, green). The near critical mode (blue) has *λ ≈* 1. **C** The variance (left axis, points along purple dashed line) and eigenvector component sum (right axis, black bars) determine how each mode contributes to the population average. The variance increases monotonically with Re(*λ*), but the eigenvector sum is independent of Re(*λ*). The critical mode has the greatest variance, and a large eigenvector sum. **D** The population sum is largely determined by the critical mode. **E** The critical mode (blue) coefficient time series is accurately revealed by the population average (red). **F-K** Avalanche size distributions (F), avalanche duration distributions (G), size versus duration (H), fluctuation functions (I), power spectra (J), and autocorrelation functions (K) are shown for the mode coefficient and population average time series shown in E. Only the near critical mode exhibits strong evidence for criticality (large power-law ranges (F, G, I, J), crackling noise scaling (H), and large autocorrelation time (K); all this evidence for criticality is clear in the population average. In panels F-H, data are shifted vertically for clarity.

### Case 2. Hidden criticality with anticorrelations

Next, we demonstrate a simple, experimentally relevant case of hidden criticality. Many experiments have reported sub-populations of neurons with strongly anti-correlated firing [20–26]. Anti-correlated activity can cancel out when averaged into the population mean, thus hiding critical dynamics [22]. Here we set up the interaction matrix *A* to obtain such anticorrelated activity.

We constructed *A* according to a commonly studied network structure with two excitatory neural populations that inhibit each other via intermediary inhibitory populations (Fig. 3A, Methods), traditionally used in studies of persistent activity and working memory [38, 39], decision making [40], and more recently in the context of criticality [22]. Like the last example, this is a case of very low-dimensional criticality – only one mode is very close to criticality with a purely real eigenvalue (a second real mode is also somewhat close with *k* ≈ 0.7, Fig. 3B). Unlike the example in Fig. 2, here the mode nearest to criticality has an eigenvector with near-zero component sum, hiding its dynamics in the population sum. Thus, analyzing the population sum reveals poor evidence for criticality. When we extract the dynamics of this hidden mode (its coefficient time series *α*(*t*)) and analyze it with traditional methods, we reveal strong evidence for criticality (Fig. 3F-K).

**FIG. 3.**
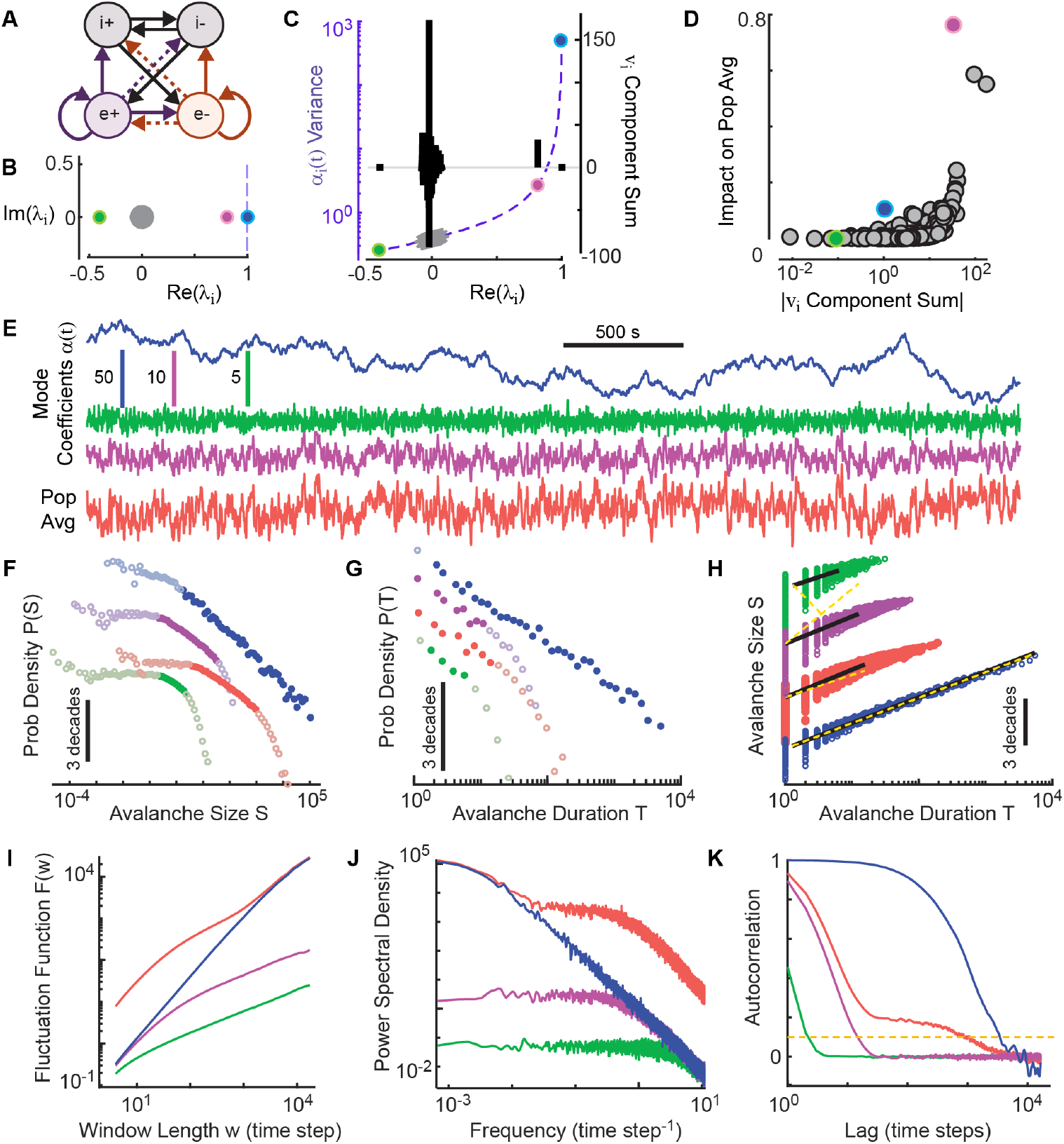
Case 2: Hidden critical dynamics due to anticorrelated activity. All panel descriptions are the same as Fig. 2; important differences are described here. **A** Schematic of interaction matrix *A* with two groups of excitatory neurons (e+ and e-) that compete via crossing inhibition. Dale’s law is enforced. **B** Bulk of eigenvalues of A are near zero (gray). The critical mode (blue) and the mode second nearest criticality (magenta) have *λ ≈* 1 and *λ ≈* 0.75, respectively. **C** The coefficient time series of the critical mode has the greatest variance, but near-zero eigenvector sum. The mode second nearest to criticality (magenta) has large variance and non-zero eigenvector sum. **D** The population average is largely determined by the mode second nearest to criticality and two modes in the bulk, but minimally impacted by the critical mode. **E** Only the near critical mode (blue) has long time scale critical dynamics in its coefficient time series. Note the strong correlation between the population average (orange) and the mode second nearest to criticality (magenta). **F-K** Only the near critical mode exhibits strong evidence for criticality (large power-law ranges (F, G, I, J), crackling noise scaling (H), and large autocorrelation time (K); this evidence for criticality is obscured in the population average.

### Case 3. Hidden criticality with oscillations

Finally, we highlight a richer case, with no single dominant mode. In this case we construct the interaction matrix according to a recent study of covariance structure of neural populations [29]. The entries of *A* are drawn from a Gaussian distribution with zero mean and then normalized to ensure the mode nearest criticality has *k* = 0.999 (Methods). The result is a mixture of multiple, coexistent near-critical modes, some of which are hidden, some with critical oscillations (Fig. 4). The population average in this case has relatively weak and messy evidence for criticality; we must analyze each mode separately to reveal strong evidence for criticality. Moreover, when analyzing the critical oscillatory mode, we must isolate the amplitude envelope before evidence for criticality becomes clear (magenta, Fig. 4F-K). If the oscillatory mode is analyzed directly without isolating the amplitude envelope, it will be dominated by the single time scale of the oscillations and confound traditional methods. This case offers a plausible explanation of traditional LRTC analysis, which also involves analyzing the envelope of particular frequency bands [8–10].

**FIG. 4.**
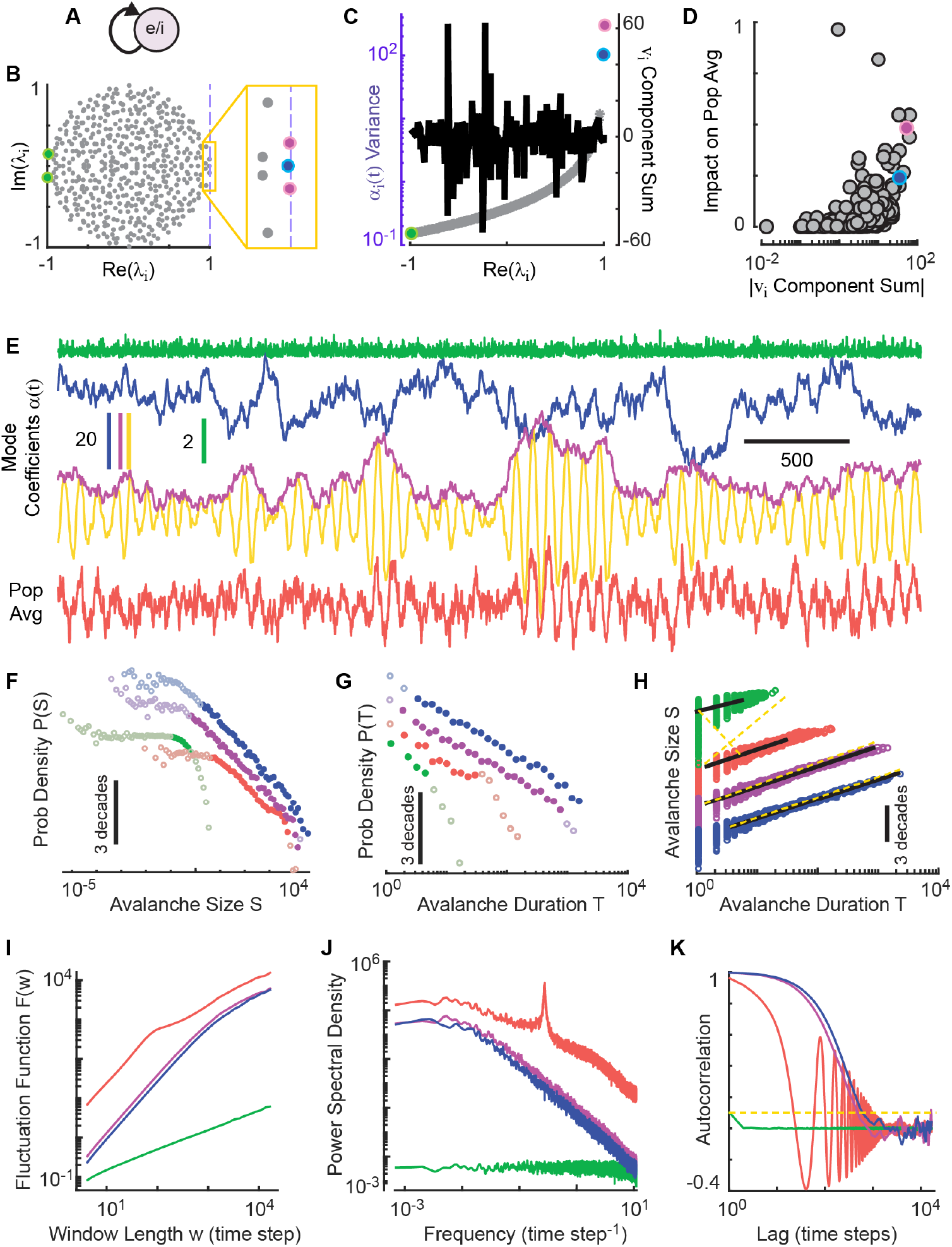
Case 3: Hidden critical dynamics in mixed modes. All panel descriptions are the same as Fig. 2; important differences are described here. **A** The interaction matrix *A* is a random matrix with approximately equal numbers of positive and negative entries drawn from a Guassian distribution. Dale’s law is violated. **B** Eigenvalues of *A* fill out the unit circle, with many modes relatively close to criticality. The mode nearest criticality is oscillatory (*λ* = 0.994 *±* 0.08*i*, magenta). The second nearest mode is non-oscillatory (*λ*=0.992, blue). **C, D** The two modes nearest to criticality have very large variance but small eigenvector sum. **D** The population average is impacted by the mode nearest to criticality, but many other modes also have substantial impact. **E** Both the near critical modes have long time scale fluctuations in their coefficient time series (blue, magenta). To reveal the critical dynamics in the oscillatory mode (*α*(*t*) - yellow), we must isolate the amplitude envelope (| *α*(*t*) | - magenta). **F-K** Critical dynamics are obscured in the population average (red) due to the mixture of modes, which is strongly impacted by the oscillations of the critical oscillatory mode. Signatures of criticality are clear for the envelope of the oscillatory mode (magenta).

In Figs. 2,3, and 4, we highlight the behavior of a few specific modes compared to the population average. For a more complete view of each case, we analyzed four properties for each individual mode and compared these properties to *d*_2_ (Fig. 5A). The four properties were chosen because they are derived from the most commonly used traditional methods of studying criticality in neural system. The first property is derived from avalanche analysis - the avalanche size distribution power-law range. Larger power law range is expected closer to criticality [7, 22]. The second property is the estimated exponent of the power spectrum, which has been reported in many previous studies [41–44]. A power-law power spectrum indicates temporal scale-invariance; an exponent near -2 is expected for directed percolation and random walks [34]. The third property is motivated by traditional LRTC analysis, which is based on DFA analysis of the power envelope of a chosen frequency band [8–10]. We estimate the exponent of the DFA fluctuation function here, which is expected to be near 1.5 for a random walk. Finally, we obtained an estimate of the longest time scales of the dynamcis from the autocorrelation function, which is also expected to be larger when closer to criticality due to critical slowing down [45, 46]. For case 1 (non-hidden criticality), these properties were nearly identical for the population sum and the dominant mode (Fig. 5B). In case 2 (anticorrelated, hidden criticality), the population sum behaved like none of the single modes, with properties that fell between those of the two modes nearest criticality (Fig. 5C). For case 3 (mixed modes, hidden criticality), the population average was also a poor representation of all single modes (Fig. 5D). Thus, we confirm that analyzing the population average is generally prone to incorrect conclusions about whether the system has any modes near criticality.

**FIG. 5.**
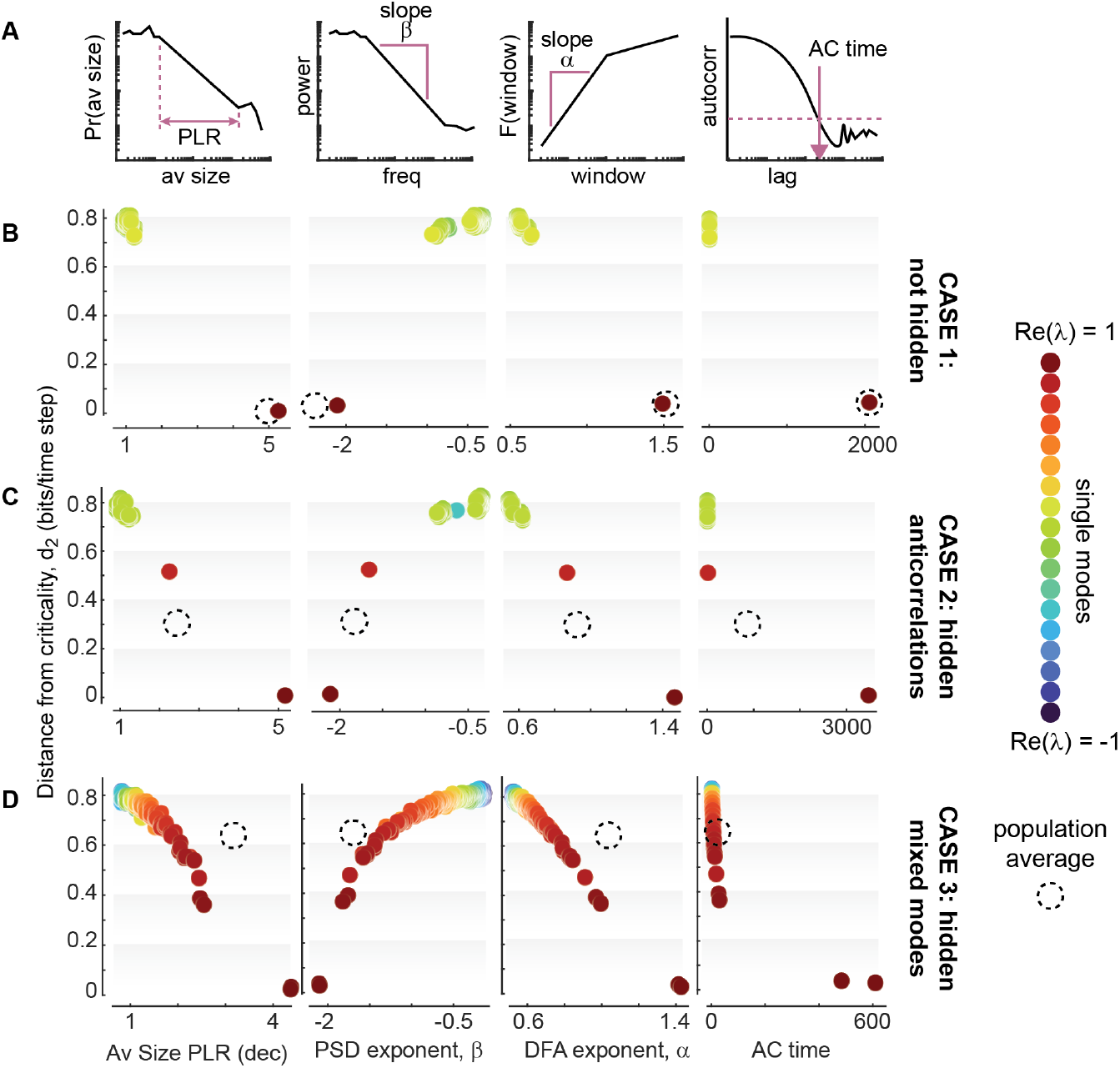
Population average often misrepresents single mode properties. **A** We analyze avalanche size power-law range, exponent of the power spectrum, DFA exponent, and autocorrelation time for each single mode. We compare these to a rigorous measure of proximity to criticality (*d*_2_). **B** For case 1, the population average correctly reveals the critical mode. **C**,**D** For cases 2 and 3, the population average obscures criticality and does not accurately represent any single mode. Color indicates *Re*(*λ*) for each single mode; dashed circles indicate the population average.

## IV. DISCUSSION

Here we report mathematical arguments and simulations, computational demonstrating how critical dynamics can be present in high-dimensional neural activity, but hidden from standard methods for assessing criticality. Any method based on a population average signal is at risk for false negative conclusions about the presence of criticality.

Several previously reported experimental results might be explained by our results. The most clear case is the anticorrelated groups of neurons reported by Jones et al [22]. The authors suggested that they had identified a new kind of criticality because most previous theoretical models did not address the case with anticorrelated neurons (Fig. 1 and Fig. 3). Our work here indicates that this is not a new kind of criticality, if the critical mode is correctly isolated, then its dynamics conform to all the traditional predictions. Similarly, Dahmen and colleagues pointed to anticorrelations as a “second type” of criticality, but did not perform traditional analyses like we did here [21]. While we agree with their point that anticorrelations can lead to hidden criticality, we would argue that all the traditionally predicted scaling laws hold for their “second type” of criticality provided that the critical modes are correctly isolated. In this sense, it is reasonable to consider all these as the same kind of criticality, but with the cautionary note: we must correctly assess each mode when many modes are mixed and potentially oscillatory.

Although most traditional methods for assessing criticality are based on the population average, a prominent exception is the phenomenological renormalization group (pRG) approach. First developed by Meshulam and colleagues [47], and subsequently adopted by several others [27, 48–50], pRG examines statistical properties of activity time series of clusters of neurons with membership to a cluster defined by how correlated the neurons are. The activity of one cluster is the average over all the neurons in the cluster - a within-cluster population average. pRG seeks power-law scaling relationships between these properties and cluster size. Because pRG groups together correlated neurons, it will naturally tend to avoid averaging anticorrelated neurons together, which may help avoid hiding criticality as we discuss here. However, as cluster size is increased, pRG ultimately considers signals that approach the population average, which will miss hidden criticality. Thus, pRG is not an ideal solution to the challenge of assessing hidden criticality.

Our results raise an important possibility. It is plausible that strong evidence for criticality is present in real data, but has been missed due to inadequate analysis methods. Many previous experimental results have shown avalanche distributions that are roughly power-law-like, but are rather messy, without a large power-law range. Our results emphasize that such results could in fact reflect partially hidden criticality, especially if some neurons exhibit strong negative correlations with others. New data analytic approaches that properly account for mixed modes and anticorrelated neurons have the potential to reveal a greater prevalence of criticality in the brain than traditional methods have indicated.

## V. METHODS

*Creation of Fig. 1 –* To simulate the single mode time series in panels A and D of Fig. 1, we approximated eq. 7 with the following discrete time equation

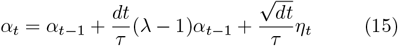

where *t* = 1, 2, …*T*, the *η*_*t*_ are drawn from a Gaussian distribution with zero mean and unit variance, *dt* = 0.001 and *τ* = 1. The basis vectors in panels B and E of Fig. 1 were constructed as follows: *v*_1,*i*_ = *i* − *N/*2 + *χ*_1,*i*_ for panel A; *v*_1,*i*_ = *i* − *N/*4 + *χ*_1,*i*_ and *v*_2,*i*_ = *N/*4 − *i* + *χ*_2,*i*_ for panel E, where *i* = 1…*N, N* = 100, and *χ*_1,*i*_ and *χ*_1,*i*_ were drawn from a Gaussian distribution with zero mean and variance of 10. The multivariate activity rasters in panels B and E were generated according to the equations just above each raster. Color indicates spike rate (blue is low, yellow is high). The spike rasters in panels C and F were created from the activity rasters rasters. Each neuron fired with probability proportional to the activity multiplied by a random constant to simulate different firing rates across neurons. The population average activity *X*(*t*) was obtained by counting spikes in time bins of duration 500*dt*.

### Impact on population average

In Figs. 2D, 3D, and 4D, we report the impact each mode has on the population average. This impact for mode *j* is defined as 1 minus the Pearson correlation coefficient of the population average *X*(*t*) and a 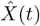, where 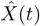 is the sum in eq. 2 excluding the *j*th term. Thus, if removing a mode results in a 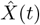 that is totally uncorrelated with *X*(*t*), then it has maximal impact of 1. If removing the mode leaves 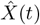 unchanged (same as *X*(*t*)), then the impact will be zero. *Multivariate rate model simulations –* To generate the data in Figs. 2, 3, and 4, we ran simulations of the following discrete time equation, which approximates eq. 3,

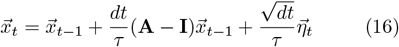

where *t* = 1, 2, …*T*, the *η*_*t*_ are drawn from a Gaussian distribution with zero mean and unit variance, *dt* = 0.002 and *τ* = 0.05. For all the results in Figs. 2, 3, and 4, we simulated *T* = 4 *×* 10^6^ time steps. To interpret model time steps in relation to experiments, We consider *τ* to be 50 ms, a characteristic time for response to input. Thus, the model time step can be interpreted as 1 ms. To better compare to typical experiments, we coarse grained in time and downsampled to a 46 ms time step, by performing a moving average in a 23 point window, with no overlap between consecutive windows.

For case 1 (Fig. 2), the network consisted of N=500 neurons, of which 20% were designated as inhibitory and the remaining 80% as excitatory. The resulting *A* matrix is sparse, with approximately 20% of its entries non-zero (corresponding to the connection probability p=0.2). The remaining 80% of the entries were set to zero, representing the absence of synaptic connections between most neuron pairs. For excitatory neurons, non-zero synaptic weights were drawn independently from a uniform distribution over the interval [0, *w*], with *w*=0.016. Inhibitory synaptic weights were drawn from [0, *gw*], where *g*=1.02, and then negated. This enforces Dale’s law; each neuron has outgoing synapses of only one sign.

For case 2 (Fig. 3), the network consisted of four neuronal populations: two excitatory groups (e+ and e-) and two inhibitory groups (i+ and i-). Each excitatory group (labeled e+ and e- in Fig. 3A) contained *N*_*e*_ = 200 neurons, and each inhibitory group contained *N*_*i*_ = 50 neurons, yielding a total of *N* = 2(*N*_*e*_ + *N*_*i*_) = 500 neurons. The full connectivity matrix *A* initialized to zero and populated probabilistically with both dense and sparse blocks. Dense connections were assigned with probability *p*_*dense*_ = 0.5, while sparse blocks were assigned with *p*_*sparse*_ = 0.05. Connections were randomly drawn independently for each eligible pair of neurons. Within each excitatory group connectivity was dense. Cross-group excitatory connectivity (between e+ and e-) was sparse. Inhibitory motifs were structured to support competition between excitatory populations (crossing inhibition): e+ projected to i-, and e- to i+, while i+ inhibited e+, and i-inhibited e-(Fig. 3A). Additionally, inhibitory neurons formed dense within-group connections. Excitatory synaptic weights were assigned a fixed value of +1, while inhibitory weights were set to -1, thus enforcing Dale’s law. Finally, the matrix was normalized to have the real part of its largest eigenvalue equal to 0.999.

For case 3 (Fig. 4), we considered dense recurrent connectivity among 500 nodes. *A* was constructed with weights drawn from a Gaussian distribution with zero mean and variance of 2. Self-connections were removed by setting all diagonal elements Aii to zero. No additional structural constraints such as symmetry or Dale’s law were imposed.

### Avalanche analysis

We performed avalanche analysis on either the coefficient time series *α*(*t*) for particular modes or on the population average time series. In both cases, avalanches were defined to be periods of time when the time series exceeded a threshold, as in many previous studies (e.g. [7, 32, 51, 52]). We chose the threshold to be the median of the time series. Avalanche size *S* was defined as the area between the time series and the threshold; avalanche duration *D* was defined as the time spent above threshold.

We followed previously developed methods for measuring power-law range for each size and duration distribution [7, 22]. In brief, this entails fitting a power-law (maximum likelihood estimation) to the avalanche distribution between a minimum *x*_*min*_ and maximum *x*_*max*_ avalanche size (or duration). All possible combinations of *x*_*min*_ and *x*_*max*_ were tried, and a goodness-of-fit was obtained for each choice of *x*_*min*_ and *x*_*max*_, where goodness-of-fit was defined as the fraction *F* of the range between *x*_*min*_ and *x*_*max*_ (in decades) for which the measured distribution fell within a 3% margin around the best fit distribution. We maximized the *x*_*min*_ to *x*_*max*_ range, subject to the constraint *F >* 0.95. The power-law range was quantified as the number of decades between *x*_*min*_ and *x*_*max*_ (*PLR* = log_10_(*x*_*max*_) −log_10_(*x*_*min*_). In Figs. 2F, 2G, 3F, 3G, 4F, and 4G, the light colored points are outside the range that is well-fit by a power-law.

To test the crackling noise scaling law, we asked whether avalanche size scales with avalanche duration according to a power-law with exponent predicted to be *γ* = (*τ*_*D*_ − 1)*/*(*τ*_*S*_ − 1), where *τ*_*D*_ is the exponent of the duration distribution and *τ*_*S*_ is the exponent of the size distribution. Both *τ*_*D*_ and *τ*_*S*_ were taken to be the exponents from the best fit power laws obtained while calculating the power-law range as described in the last paragraph. To test whether crackling noise scaling holds, we compared *γ* to the slope of a best fit line to the log(S) vs log(D) relationship. The slopes of the yellow dashed lines in Figs. 2H, 3H, and 4H are *γ*; the slopes of teh black solid lines in Figs. 2H, 3H, and 4H are the best fit slope. Mismatch between the yellow dashed line and the black line indicates that crackling noise scaling does not hold.

### DFA analysis

We considered 10 window sizes per decade, logarithmically spaced, between 4 points and one tenth the time series duration. For a window duration *w*, we broke the time series into consecutive segments (with 50% overlap), each segment with duration *w*. Each segment was detrended (a best fit linear trend was subtracted) and then its standard deviation was calculated. The fluctuation functions in Figs. 2I, 3I, and 4I, report the average standard deviation versus the window duration *w*. To mitigate finite-size effects, we restricted the regression to the lower 70% scales in log space (i.e., excluded the largest 30% of *w*. The DFA scaling exponent *α* was then obtained as the ordinary least-squares slope of the log-log relation log_10_ *F*(*w*) = *c* + *α* log_10_ *w*, using the retained scales.

### Power spectra

We computed the power spectral density using Welch’s method (Matlab’s *pwelch* function). We used windows of duration one tenth the full time series duration. Each window overlapped the next by 50%. To avoid low-frequency bias, we restricted the fit to frequencies *f* ≥ _*cut*_, where *f*_*cut*_ is the 30th percentile of the logarithmic frequency grid between the minimum positive sampled frequency and the maximum frequency. We then performed an ordinary least squares regression in log-log coordinates and reported the fitted slope *β* as the spectral exponent.

### Autocorrelation time estimate

We quantify a characteristic time for temporal correlations by identifying the smallest lag for which the autocorrelation function falls below a threshold of 0.1. These time scales were assessed for each mode in Fig. 5B, C, and D and reported as “AC time”.

## Notes

### Competing Interest Statement

The authors have declared no competing interest.

### Funding

- National Institutes of Health: R15NS135396, R01DA060744

